# Survival Beyond the Perinatal Period Expands the Phenotypes Caused by Mutations in *GLE1*

**DOI:** 10.1101/124172

**Authors:** Edith Said, Jessica X. Chong, Maja Hempel, Jonas Denecke, Paul Soler, Tim Strom, Deborah A. Nickerson, Christian Kubisch, Michael Bamshad, University of Washington Center for Mendelian Genomics, Davor Lessel

## Abstract

Mutations in *GLE1* underlie Lethal Congenital Contracture syndrome (LCCS1) and Lethal Arthrogryposis with Anterior Horn Cell Disease (LAAHD). Both LCCS1 and LAAHD are characterized by reduced fetal movements, congenital contractures, and a severe form of motor neuron disease that results in fetal death or death in the perinatal period, respectively. Via trio-exome sequencing, we identified bi-allelic mutations in *GLE1* in two unrelated individuals with motor delays, feeding difficulties and respiratory insufficiency who survived beyond the perinatal period. Each affected child had missense variants predicted to result in amino acid substitutions near the C-terminus of GLE1 that are predicted to disrupt protein-protein interaction or GLE1 protein targeting. We hypothesize that mutations that preserve function of the coiled-coil domain of GLE1 cause LAAHD whereas mutations that abolish the function of the coiled-coil domain cause LCCS1. The phenotype of LAAHD is now expanded to include multiple individuals surviving into childhood suggesting that LAAHD is a misnomer and should be re-named Arthrogryposis with Anterior Horn Cell Disease (AAHD). Too few cases have been reported to identify significant genotype-phenotype relationships, but given that perinatal lethality in AAHD typically resulted from respiratory failure, it is possible that early or aggressive airway management such as early tracheostomy and ventilation may enable survival beyond the perinatal period.

## INTRODUCTION

Whole exome sequencing (WES) is a powerful tool for dissecting the genetic basis of Mendelian conditions, including both the identification of novel disease genes and the diagnosis of monogenic diseases (Chong et al., 2015a). WES has been particularly useful for making a precise molecular diagnosis of phenotypically and genetically heterogeneous disorders. In turn, this has broadened the spectrum of phenotypes of many known Mendelian conditions and the number of conditions associated to single genes (Dyment et al., 2015; Thevenon et al., 2016).

Mutations in *GLE1* have previously been reported to underlie two autosomal recessive conditions, Lethal Congenital Contracture Syndrome 1 (LCCS1; OMIM #253310) and Lethal Arthrogryposis with Anterior Horn Cell Disease (LAAHD; OMIM #611890), which result in death in the fetal and perinatal period, respectively (Nousiainen et al., 2008). LCCS1 is characterized by a lack of fetal movements in the second trimester of pregnancy. Affected individuals typically present with intrauterine growth retardation (IUGR), fetal hydrops, micrognathia, pulmonary hypoplasia, and multiple joint contractures (Herva et al., 1985). Individuals with LAAHD present with a similar albeit “milder” phenotype, with prenatal onset of diminished fetal mobility and contractures, and post-natal respiratory failure resulting in perinatal death (Vuopala and Herva, 1994). IUGR and hydrops fetalis are either absent or mild in LAAHD. Pathological analysis of persons with LCCS1 reveals a lack of anterior horn motor neurons, atrophy of the ventral spinal cord and extreme skeletal muscle atrophy (Herva et al., 1985). Infants with LAAHD demonstrate varied abnormalities of skeletal muscle ranging from minor variation in fiber diameter to groups of atrophic fibers with hypertrophic type I fibers, and anterior horn cell neurons being degenerated or diminished (Vuopala and Herva, 1994).

Herein we report the discovery of two unrelated individuals with a phenotype similar to LAAHD, who survived beyond 4 years of life, and were either homozygous or compound heterozygous for missense variants in *GLE1*. Each affected child had global developmental delay, severe motor abnormalities and respiratory insufficiency. These findings expand the phenotype of LAAHD and suggest that either some mutations in *GLE1* are associated with better outcomes and / or differences in intervention early in life can improve outcomes.

## MATERIAL AND METHODS

### Human Subjects

All biological samples and photographs were obtained following written informed consent from affected individuals or their legal representatives. The study was approved by the University Medical Center Hamburg-Eppendorf, Malta, and the University of Washington (Seattle, USA). The study was performed in accordance with the Declaration of Helsinki protocols. Investigators studying the two affected individuals described here connected via GeneMatcher (Sobreira et al., 2015), a member of the MatchMaker Exchange (MME). MME is a network of web-based tools for researchers and clinicians to share candidate genes (Philippakis et al., 2015).

### Exome sequencing and variant validation

Exome sequencing of DNA samples from both parents in Family A and proband 1 was performed on exons captured by Roche Nimblegen SeqCap EZ Human Exome Library v2.0 and sequenced using the Illumina HiSeq2500/4000 as previously described (Chong et al., 2015b). Exome sequencing of DNA samples from both parents in Family B and proband 2 was performed on exons captured by SureSelect Human All Exon 50Mb V5 Kit (Agilent, Santa Clara, CA, USA) and sequenced using a HiSeq2500 (Illumina, San Diego, CA, USA) as described before (Hempel et al., 2015). Candidate variants were validated and segregation confirmed by Sanger sequencing of DNA from all informative and available family members as previously described, Family A (Chong et al., 2015b) and Family B (Lessel et al., 2015).

## RESULTS

### Family A

Proband 1 (Figure 1; Table I) was the first-born male of healthy non-consanguineous parents of European / Maltese ancestry. The pregnancy was complicated by decreased fetal movements and polyhydramnios. Due to fetal bradycardia with initial uterine contractions he was delivered by emergency Caesarian section at 38 weeks gestation with a birth weight of 2,480 g (-2.0 SD), a birth length of 46 cm (-2.3 SD) and a ocipitofrontal head circumference (OFC) of 35 cm (0 SD). APGAR scores were 5, 6 and 7 at 1^st^, 5^th^ and 10^th^ minutes, respectively. Severe respiratory distress required resuscitation and admission to the neonatal intensive care unit where he was ventilated. Abnormal facial characteristics included a prominent forehead, down-slanting palpebral fissures, tent-shaped mouth, prominent frenulum, micrognatia and low-set ears (Figure 1A). Congenital contratures included bilateral camptodactyly of both hands, adducted thumbs, and bilateral clubfeet. His neurological exam at birth was notable for hypertonia of his upper and lower limbs with startling and jerky movements. Oropharyngeal secretions were excessive and due to swallowing difficulties, he required nasogastric feeding.

He was eventually weaned off the ventilator but required oxygen by continuous positive airway pressure intermittently, especially at night. A sleep study showed short episodes of apnoea. Laryngoscopy showed an elliptical epiglottis with floppy larynx.

**Fig 1.**
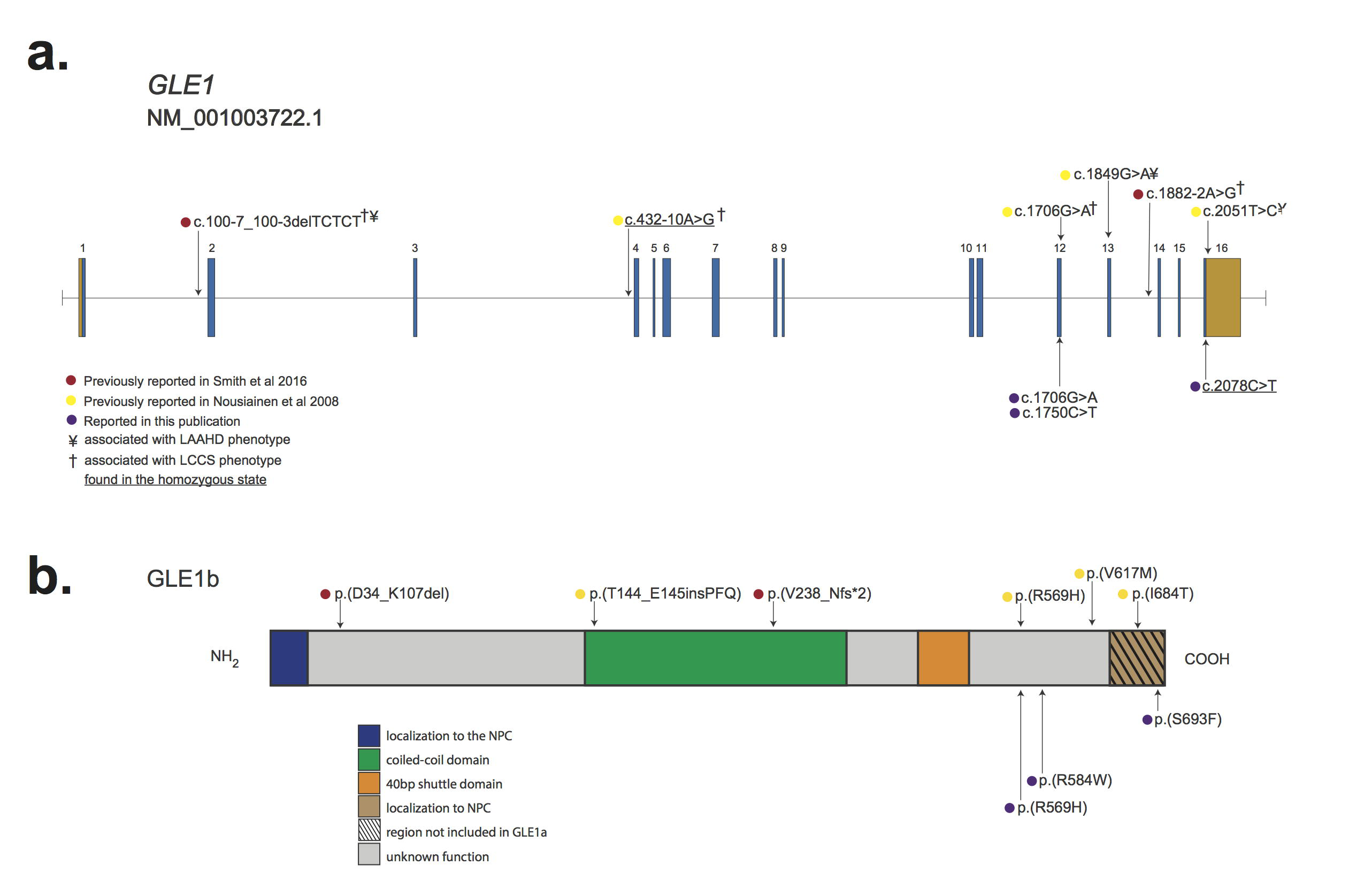
Photographs of individuals with LAAHD. (Due to biorxiv.org rules, photographs have been omitted from this preprint) Images show facial features of proband 1 at 8 weeks (A-1) and 9 months (A-2, A-3) and proband 2 (B-1, B-2) at 3 years 5 months of age. Features include prominent forehead, downslanting palpebral fissures, and micrognathia. See Table I for detailed clinical information on each affected individual.

**Table I.**
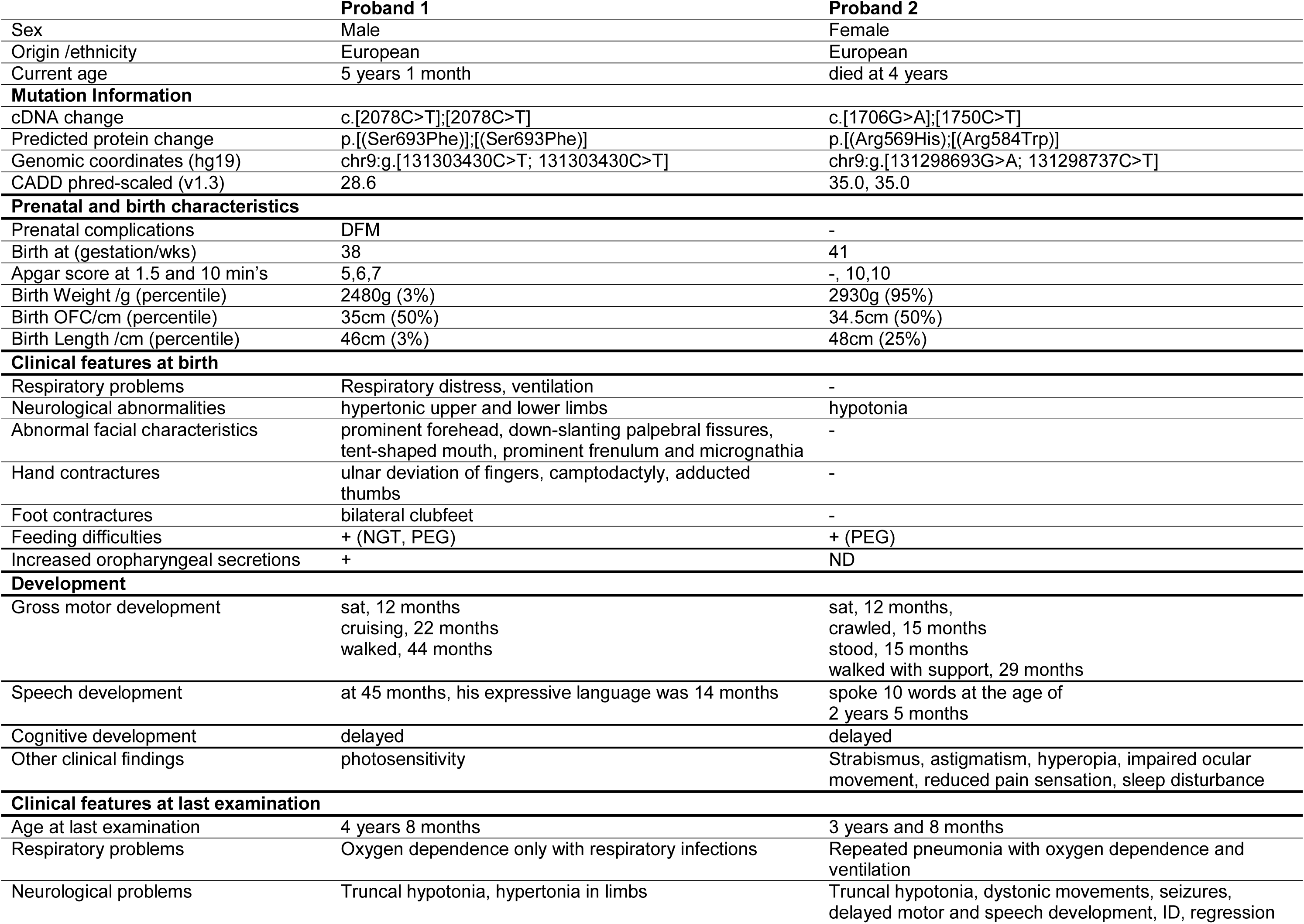
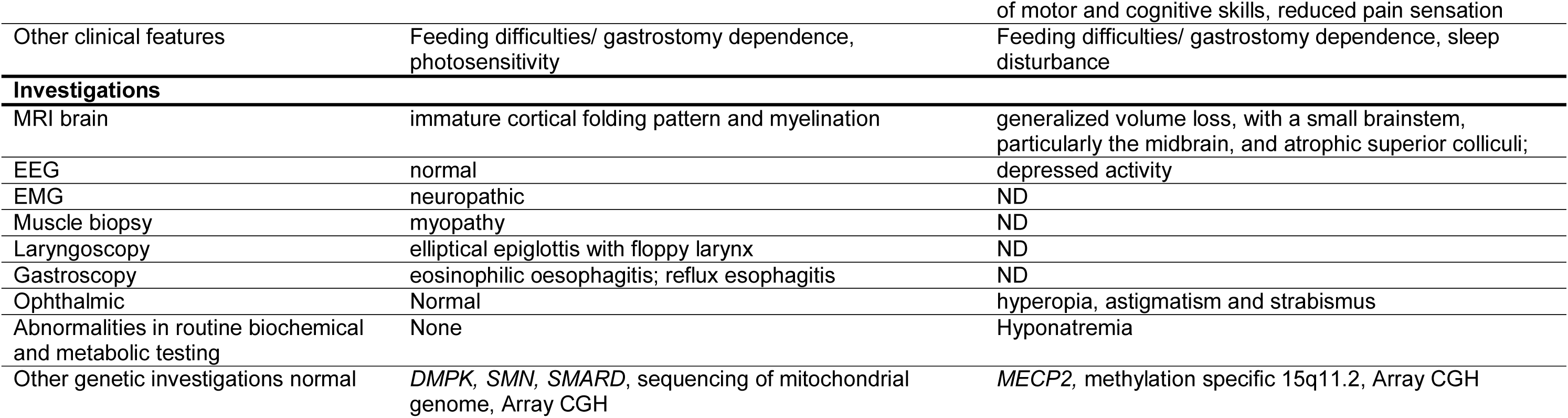
Mutations and Clinical Findings of Individuals with LAAHD. This table provides a summary of clinical features and clinical investigations of affected individuals in this study. Plus (+) indicates presence of a finding, minus (-) indicates absence of a finding, ND = no data were available. N/A = not applicable. CADD = Combined Annotation Dependent Depletion. DFM = decreased fetal movements. cDNA positions provided as named by the HGVS MutNomen web tool relative to NM_001003722.1.

At the age of 12 months, he was not feeding adequately and a percutaneous endoscopic gastrostomy tube was placed. His muscle tone was variable with brisk reflexes. He had a flexion contracture of the left hip, decreased abduction of both shoulders and decreased extension of both elbows. Bilateral ulnar deviation of the hands and thumb adduction were still evident. An EMG showed motor neuropathic and a muscle biopsy non-specific myopathic features. Analysis of respiratory chain enzymes in muscle was normal. Brain MRI demonstrated immature myelination and cortical folding but was otherwise normal. Echocardiography, a creatine kinase level and extensive metabolic studies were normal. Results of arrayCGH and sequencing of *DMPK, SMN1* and *UBA1* did not identify any pathogenic variants.

His motor and speech milestones were delayed. He sat unsupported at 10 months and walked with a broad base gait at 3 years 8 months. At age of 18 months he could say a few words, and at 3 years 9 months, his expressive language was estimated to be that of a 10 to 14 month-old child. Cognitive testing revealed a development level comparable with a 20 to 24 month-old child. Examination at 4 years 8 months of age revealed he still had an expressive speech delay due mainly to his micrognathia but his receptive language was normal. He had truncal hypotonia with brisk reflexes at the knees and ankles. His muscle bulk was markedly diminished. He could walk without assistance but occasionally used a K-walker. He had not needed supplemental oxygen for 18 months. He still had difficulty in swallowing solids and was fed on semisolids with supplements by gastrostomy tube.

### Family B

Proband 2 (Figure 2; Table I) was the first-born female of healthy non-consanguineous parents of European ancestry. A previous pregnancy was terminated at 20 weeks because of generalized non-immune hydrops fetalis, however no genetic analysis was performed and fetal DNA was not obtained. After an uneventful pregnancy, Proband 2 was born at 41 weeks gestational age via vacuum extraction due to fetal bradycardia. Her birth weight was 2,930 g (- 1.4SD), birth length 48 cm (-1.8SD) and OFC 34.5 cm (-0.4SD). APGAR scores were 10 at both 5 and 10 minutes. Family history was unremarkable for developmental delay, neurological or immunological problems.

**Fig 2.** Genomic and protein structure of GLE1 and spectrum of pathogenic mutations in *GLE1*. A) *GLE1* is composed of 16 exons, all which are protein-coding (blue) and two of which also contain non-coding sequence (orange). Lines with attached dots indicate the approximate locations of eleven different recessive variants that cause LCCS1 and LAAHD. The color of each dot reflects the report of each mutation. B) Protein topology of GLE is comprised of six domains. Most mutations lie in the C-terminal portion of GLE1. No mutations in the N-terminal Nup155 binding domain or in the nucleocytoplasmic shuttling domain have been reported. The approximate positions of variants that cause LCCS1 and LAAHD are indicated by colored dots. Only p.(V238_Nfs*2) has been shown to result in complete lack of GLE1; all other variants have been shown or are expected to be expressed.

She was evaluated at 12 months of age because of suspected developmental delay. Her parents had noted feeding difficulties, decreased spontaneous movements and hypotonia since birth. She suffered from sleep-onset insomnia which was treated with melatonin. Developmental milestones were delayed. She sat unsupported at age of 12 months, crawled at 15 months and stood at 15 months. She walked with support at the age of 2 years and 5 months. Her speech development was also delayed. A 2 years and 5 months, she could speak ten words, but not a full sentence. Parents reported decreased sensitivity to pain. Starting at the age of 14 months she developed frequent febrile infections. These infections were accompanied by regressions in development. Each infection resulted in loss of motor skills and a remarkable extended recovery time without a return to baseline function. At the age of 3 years 3 months, she experienced a first febrile seizure.

Examination at age of 3 years and 3 months (Figure 1B) revealed a restless girl with truncal hypotonia, dystonic movements of the upper limbs, involuntary facial grimacing and eye movements, and unmotivated yelling. She was unable to sit or walk unsupported, and protective reflexes were depressed. Supported gait was ataxic. Her weight, height and OFC were within the normal range. Basic metabolic studies revealed severe hyponatremia. (121 mmol/L, normal value 135 – 145 mmol/L). Extensive metabolic testing, including serum and CSF lactate were normal. MRI of brain showed slight atrophy, but no signal alteration in basal ganglia or lactate peak in spectroscopy. EEG revealed deceleration of basic activity. Eye examination showed hyperopia, astigmatism and strabismus.

At the age 3 years 8 months, she had parainfluenza pneumonia which progressed to severe hypoxemic respiratory failure requiring ventilation. Hyponatremia (126 mmol/L) was observed as in the previous admissions. After prolonged weaning from the ventilator, she developed delirium and motor abnormalities with flailing of arms and legs, grimacing and writing movements, even when she was asleep. These involuntary movements improved over the next weeks but she had lost her gross motor skills including sitting, crawling. Repeated MRI of brain showed generalized parenchymal volume loss and a small brainstem, particularly the midbrain. A G-tube was installed for feeding. At 3 years and 11 months she experienced her first generalized seizure without fever. After her fourth bout of pneumonia she died because of respiratory insufficiency at the age of four years. Genetic testing including methylation specific MLPA in 15q11.2, *MECP2* sequencing, MLPA and array-CGH was normal.

### Trio whole exome sequencing

#### Family A

WES data were annotated with the Variant Effect Predictor v83 (McLaren et al., 2016) and analyzed with GEMINI 0.18.1 (Paila et al., 2013). Variants unlikely to impact protein-coding sequence (for which GEMINI impact_severity = LOW), variants flagged by the Genome Analysis Toolkit (GATK) as low quality, and variants with an alternative allele frequency > 0.005 in any superpopulation in EVS/ESP6500, 1000 Genomes (phase 3 release), or the ExAC Browser (v.0.3) were excluded. Variants that were frequent (alternative allele frequency >0.5) in an internal database of >2,900 individuals (Geno2MP v1.2 release) were also excluded. Analysis under different inheritance models (i.e. autosomal de novo, homozygous recessive, compound heterozygous, X-linked de novo, X-linked recessive models) identified 12 candidate genes (*de novo: SYNE1;* homozygous recessive: *GLE1, DDX12P, ODF2, C9orf9;* compound heterozygous: *CAPN8, KIRREL, FSCN2, COL6A6;* X-linked recessive: *RGAG1, USP26, DKC1*). The homozygous variant in *GLE1* (GenBank: NM_001003722.1), c.2078C>T [p.(Ser693Phe)], was considered the best candidate given the gene’s known function and the similarity of phenotypes previously reported to be caused by mutations in *GLE1*.

#### Family B

Bioinformatic filtering did not detect *de novo* variants. However, filtering for rare candidate variants (minor allele frequency MAF < 0.01, according to ExAC browser) under an autosomal recessive model of inheritance identified compound heterozygous alterations, c.1706G>A [p.(Arg569His)] and c.1750C>T [p.(Arg584Trp)] in only a single gene, *GLE1*.

### Validation and segregation analysis

Sanger sequencing confirmed that case 1 was homozygous and both parents were heterozygous for the *GLE1* variant c.2078C>T [p.(Ser693Phe)]. Even though the parents were non-consanguineous, the proband’s homozygosity for a single variant and the parents’ origin from the same island suggested that c.2078C>T [p.(Ser693Phe)] might be a founder mutation.

Indeed, the variant was found within a large 7.7 Mb run of homozygosity. Sanger sequencing confirmed both heterozygous alterations in Case 2, whereas her parents had only one variant in a heterozygous state each.

The *GLE1* variant c.2078C>T, p.(Ser693Phe) in proband 1 has not been observed in over 76,000 individuals according to publicly available databases: the Exome Variant Server (v2), 1000 Genomes phase 3, the Exome Aggregation Consortium (ExAC) browser (v0.3), or UK10K (February 15, 2016 release). The variant c.1706G>A, p.(Arg569His) identified in proband 2, has been reported before in individuals with a LCCS1/LAAHD-like phenotype (Nousiainen et al., 2008; Ellard et al., 2014) and has a global allele frequency of 0.0004 according to ExAC. The other heterozygous variant identified in proband 2, c.1750C>T, p.(Arg584Trp) has a global allele frequency of 5x10^-5^. All three variants have high phred-scaled CADD (v1.3) scores (which take into account conservation and the scores from other pathogenicity predictors such as PolyPhen-2 and SIFT), consistent with pathogenic recessive variants: 35.0 for p.(Arg569His); 35.0 for p.(Arg584Trp); and 28.6 for p.(Ser693Phe) (Table II). Taken together, both the genetic and phenotypic data suggest that the GLE1 variants identified (Table I; Figure 2) are causal for a phenotype overlapping LAAHD and LCCS1.

**Table II.** Summary of Mutations and Selected Clinical Findings in all individuals with *GLE1* mutations reported to date. This table summarizes the combinations of mutations, domains and isoforms altered by each mutation, and clinical features distinguishing LCCS1 and LAAHD in all affected individuals with *GLE1* mutations reported to date. ND = no data. NMD = nonsense-mediated decay. cDNA positions provided as named by the Mutalyzer.nl web tool relative to NM_001003722.1. Data in columns 1-4.

|  |  |  |  |  |  |  |  |  |
| --- | --- | --- | --- | --- | --- | --- | --- | --- |
| # individuals | 51 | 1 | 6 | 6 | 2 | 2 | 1 | 1 |
| # families | 28 | 1 | ND | ND | 1 | 1 | 1 | 1 |
| **Mutation 1** |  |  |  |  |  |  |  |  |
| cDNA change | c.432-10A>G | c.432-10A>G | c.432-10A>G | c.432-10A>G | c.1706G>A | c.100-7_100-3d eITCTCT | c.2078C>T | c.1706G>A |
| Predicted protein change | p.T144_E145ins PFQ | p.T144_E145ins PFQ | p.T144_E145ins PFQ | p.T144_E145ins PFQ | p.R569H | p.D34_K107del | p.Ser693Phe | p.R569H |
| Domain Function | oligomerization | oligomerization | oligomerization | oligomerization | IP6 binding | unknown | NPC localization | NPC localization |
| Isoforms affected | A, B | A, B | A, B | A, B | A, B | A,B | B | A, B |
| **Mutation 2** |  |  |  |  |  |  |  |  |
| cDNA change | c.432-10A>G | c.1706G4A | c.1849G>A | c.2051T>C | c.1849G>A | c.1882-2A>G | c.2078C>T | c.1750C>T |
| Predicted protein change | p.T144_E145ins PFQ | p.R569H | p.V617M | p.I684T | p.V617M | p.V238_Nfs*2 | p.S693F | p.R584W |
| Isoforms affected | A, B | A, B | A, B | B | A, B | A (NMD), B (NMD) | B | A, B |
| **Diagnosis** | LCCS1 | LCCS1 | LAAHD | LAAHD | "fetal akinesia" | "GLE1-associated AMC" |  |  |
| Prenatal signs | no fetal movements | no fetal movements | decreased fetal movements | decreased fetal movements | ND | ND | decreased fetal movements | "uneventful pregnancy" |
| Age | died <32 weeks gestation | termination @ 14 weeks | some live births | some live births; one child died @ 2 months | termination | 1st child died @ 14 days<br>Sibling alive @ 12 yrs | alive @ 4 yrs 8 months | died @ 4 years |
| Publication | Nousiainen et al. 2008 | Nousiainen et al. 2008 | Nousiainen et al. 2008 | Nousiainen et al. 2008 | Ellard et al. 2015 | Smith et al. 2016 | this study | this study |

## DISCUSSION

We identified two families without known Finnish ancestry that each had a child who had a phenotype similar to LAAHD and LCCS1 and was either compound heterozygous or homozygous for missense variants in *GLE1* predicted to be deleterious. In contrast to LAADH and LCCS1, both children survived the perinatal period and one is still alive (Table II). Indeed, no fetus with LCCS1 surviving beyond 35 weeks EGA or infant with LAADH surviving beyond eight weeks after birth has been reported until the recent publication of single exception (Smith et al., 2016). Accordingly, the natural history of the children reported herein is considerably different from that of children with either LCCS1 or LAADH (Table II). This observation suggests that either the children reported herein have a condition distinct from both LCCS1 and LAADH, that mutations in *GLE1* produce a single condition with a broad distribution of phenotypes, or that these children represent phenotypic expansion of LCCS1 or LAAHD.

Each of the variants we identified affects a residue(s) located in the C terminus of GLE1 (Figure 2; Table II), suggesting a possible relationship between affected domain(s) or affected isoform(s) and phenotype. However, other than proband A, who is homozygous for p.(S693F), a variant that lies in the domain unique to GLE1B, all individuals with *GLE1* mutations reported to date have either one or two mutations predicted to affect both isoforms (Table II). This makes it difficult to distinguish whether there is a relationship between observed phenotype features, severity, and type or location of a variant.

Smith et al. (Smith et al., 2016) recently reported a family with two affected brothers who had markedly different phenotypes from one another but were compound heterozygotes for *GLE1* variants, c.100-7_100-3delTCTCT [p.(D34_K107del)] and c.1882-2A>G [p.(V238_Nfs*2)]. Specifically, one brother died at two weeks of age while the other survived to 12 years. Smith et al. (Smith et al., 2016) proposed that LCCS1 and LAAHD constitute the same clinical entity with variable severity because the two disorders have “nearly indistinguishable phenotypes.” However, the observation that all individuals homozygous for p.T144_E145insPFQ died prenatally and exhibit no fetal movements during gestation (Table II) suggests that LCCS1 represents a distinct clinical entity as the prognosis is different than for individuals who are compound heterozygous or are homozygous for other *GLE1* mutations. This is similar to other human genes in which a specific mutation or at most several mutations result in a distinct phenotype (e.g., *FGFR3* and achondroplasia).

GLE1 is an essential mRNA export factor^15^ and appears to also play a role in both initiation and termination of protein translation in eukaryotes^1614^. *GLE1* encodes at least two different isoforms, GLE1A and GLE1B, with GLE1B being ∼1,000-fold more abundant across all tissues (Kendirgi et al., 2003). The two isoforms are nearly identical—specifically, both GLE1A and GLE1B contain a N-terminal domain that is required for localization to the nuclear pore complex (NPC), a nucleocytoplasmic shuttling domain, and a coiled-coil domain that is necessary for oligomerization of GLE1—oligomerization is needed for NPC localization and mRNA shuttling. The only domain distinguishing the two isoforms is a C-terminal 43 amino acid segment unique to GLE1B (Figure 2B) that is also required for localization of GLE1 to the nuclear pore complex (NPC).^2122^ Nevertheless, this difference leads to distinct functions for each isoform: GLE1B exports mRNAs from the nucleus to the cytoplasm via the NPC, while GLE1A localizes to the cytoplasm and is required for stress granule assembly and disassembly, thereby regulating protein translation (Aditi et al., 2015).

We predict that biallelic mutations that truncate or otherwise result in loss of function of the coiled-coil domain of both GLE1 isoforms cause LCCS1. In contrast, individuals with mutations that preserve the function of the coiled-coil domain in both isoforms cause LAAHD. We also suggest that because the phenotype of LAAHD is now expanded to include multiple individuals surviving into childhood, LAAHD should be re-named Arthrogryposis with Anterior Horn Cell Disease (AAHD). Perinatal lethality in AAHD typically resulted from respiratory failure, and differences in differences in clinical management such as early tracheostomy and ventilation may enable survival beyond the perinatal period.

## Acknowledgments

The authors are thankful to affected individuals and their family members for participation. University of Washington Center for Mendelian Genomics (UW-CMG) was funded by the National Human Genome Research Institute and the National Heart, Lung and Blood Institute grant U54HG006493 to Drs. Debbie Nickerson, Michael Bamshad, and Suzanne Leal. The authors would like to thank the Exome Aggregation Consortium and the groups that provided exome variant data for comparison. A full list of contributing groups can be found at
http://exac.broadinstitute.org/about. The authors would like to thank the University of Washington Center for Mendelian Genomics and all contributors to Geno2MP for use of data included in Geno2MP. The authors would like to thank the NHLBI GO Exome Sequencing Project and its ongoing studies which produced and provided exome variant calls for comparison: the Lung GO Sequencing Project (HL-102923), the WHI Sequencing Project (HL- 102924), the Broad GO Sequencing Project (HL-102925), the Seattle GO Sequencing Project (HL-102926) and the Heart GO Sequencing Project (HL-103010).

